# The DNA damaging properties of the experimental G-quadruplex (G4) drug QN-302 are potentiated by the DNA repair inhibitor Olaparib and mitigated by the molecular helicase PhpC

**DOI:** 10.64898/2026.02.06.704351

**Authors:** Garance Psalmon, Angélique Pipier, Manon Barbotte, Robert H. E. Hudson, Stephen Neidle, David Monchaud

## Abstract

**Background:** QN-302 is a tetra-substituted naphthalene diimide (NDI) compound designed to interact with G-quadruplex (G4) DNA. QN-302 is currently being evaluated in a phase 1 clinical trial on patients with advanced pancreatic ductal adenocarcinoma (PDAC) and other solid tumors. However, the mechanistic origin(s) of its anticancer activity remains to be fully understood.

**Results:** We report herein the ability of QN-302 to damage DNA at G4 sites in cancer cells. To this end, we implemented a series of *in vitro* assays (FQA and FRET-melting) and cell-based techniques (*in situ* click imaging and immunodetection) that concurred in demonstrating both the DNA damaging properties of QN-302 and its ability to engage G4s in human cancer cells. Then, we investigate its anticancer effects in PDAC (MIA PaCa-2 cells) and show that it can be efficiently potentiated upon combination with Olaparib, an inhibitor of DNA repair, in an approach referred to as chemically induced synthetic lethality.

**Conclusion:** This study not only confirms the excellent anticancer properties of QN-302 in human cancer cells but also provides insights into its mechanism of action. The optimization of this therapeutic activity by combination with Olaparib opens a promising new avenue for improving its clinical efficacy.

## Background

DNA and RNA G-quadruplexes (G4s) belong to the family of alternative nucleic acid structures whose cellular roles and functions are currently being scrutinized with a great deal of attention (1-3). The involvement of G4s is currently being studied in almost every cellular pathway (pertaining to cellular growth, differentiation and division, gene expression, genome stability, epigenomic modifications, etc.) (1, 4-6), making them new genetic switches that could be actuated to seize control over the corresponding cellular transactions. By definition, G4s can fold from guanine (G)-rich sequences (overly abundant in the human genome (7-9) and transcriptome) (10-13) in a transient manner only, when the sequence involved is freed from its duplex constraint. Low molecular weight compounds can be used to modulate G4 folding and this intervention has consequences genome-/transcriptome-wide since many G4-containing genes/transcripts and related pathways may be affected.

G4-stabilizing molecules (or G4 ligands) (14-16) have found applications in oncology (15, 17), neurobiology (18) and virology (19). Most of the efforts have been invested in the use of G4 ligands as anticancer agents and to date, only two compounds have reached clinical trials (Fig. 1A), Pidnarulex (*a*.*k*.*a*. CX-5461) (20) and QN-302 (previously known as SOP1812) (21, 22). The former is currently being assessed in cancer patients with DNA repair deficiencies (notably, defective homologous recombination (HR) pathways; see NCT02719977 (23) and NCT04890613, ongoing) while the latter is enrolled in phase 1 clinical trial on advanced pancreatic ductal adenocarcinoma (PDAC) patients (see NCT06086522).

**Figure 1:**
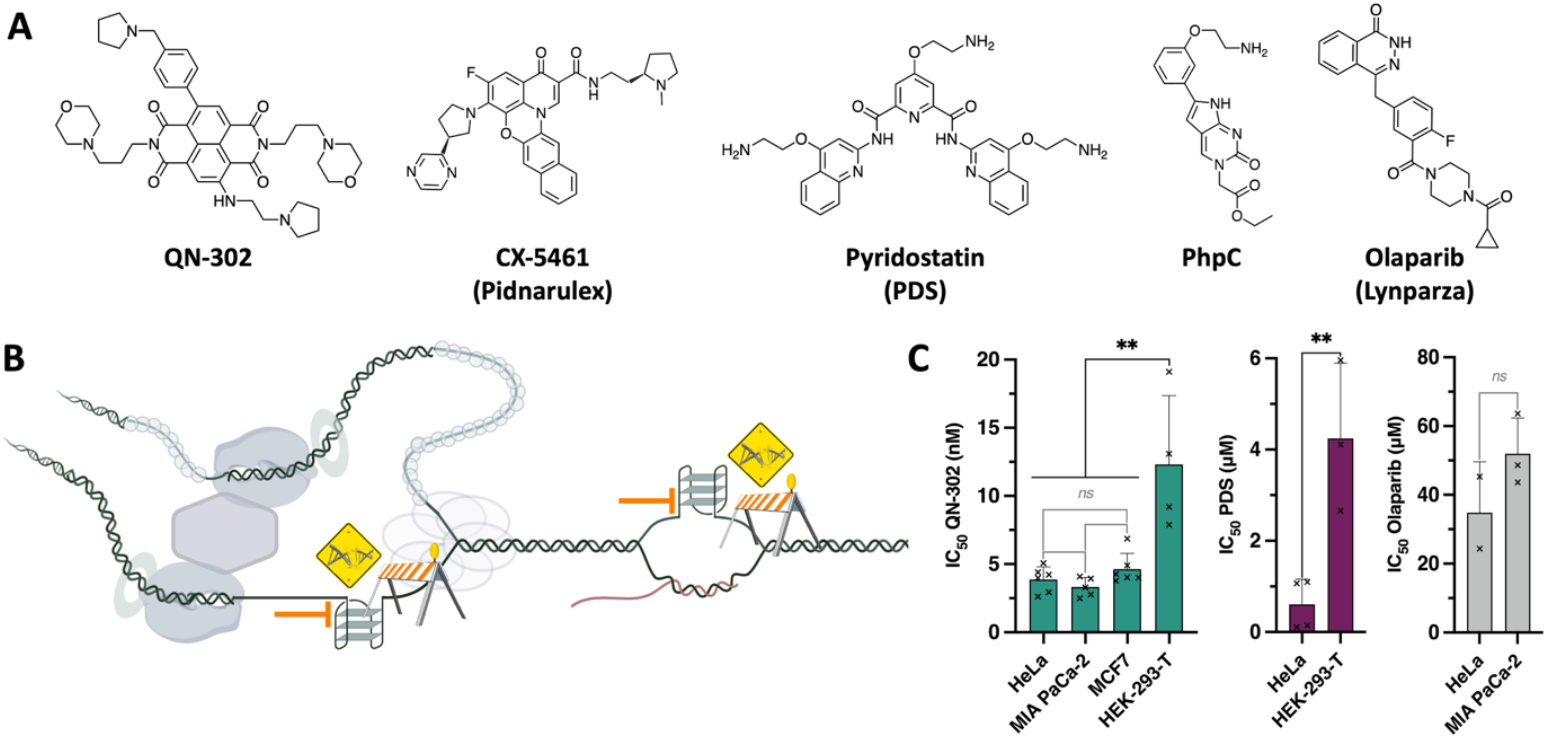
**A**. Chemical structures of G4 ligands (QN-302, CX-5461 and PDS), G4 destabilizer (PhpC) and DNA repair inhibitor (Olaparib) used and/or discussed in this study. **B**. Schematic representation of the way G4s jeopardize genetic instability by acting as roadblocks to DNA transactions, which triggers genomic instability. **C**. Toxicity of the molecules (after 72-h treatment) used in this study, established by the SRB assay, in cancer (HeLa, MIA PaCa-2 and MCF-7) and immortalized cells (HEK-293-T). Multiple unpaired t-test was employed for the statistical analyses with *: *P* ≤ 0.05, **: *P* ≤ 0.01, “ns” for non-significant: *P* > 0.05.

QN-302 is a pan-G4 ligand since it interacts with many cellular G4-forming sequences, including those found in the promoter regions of genes such as *GLI1, S100P, MAPK11, CX3CL1 and PRDM16* (24). A key feature of G4 ligands is their ability to inflict severe injuries to the genome, notably triggering G4-associated DNA damage (Fig. 1B) (25, 26). We thus wondered whether QN-302 does induce DNA damage, in a manner that could be reminiscent of what has been established with CX-5461 (20, 27, 28) and also with the reference ligand pyridostatin (or PDS, Fig. 1A) (29-31) another well-known pan-G4 ligand. To this end, we first characterize here the ability of QN-302 and PDS (used as a control) (32, 33) to engage G4s in a model of human cancer (HeLa cells) by *in situ* click imaging. Next, we show that this stabilization induces genetic instability, triggering an elevated level of double strand DNA breaks (DSBs). Quite uniquely, both G4 engagement and G4-mediated DNA damage were confirmed here by using the molecular helicase (or G4-destabilizer) PhpC (34) to resolve cellular G4s and thus, mitigate the cellular effects of G4 ligands (Fig. 1A). Finally, given that QN-302 is currently being clinically assessed (*vide supra*), we confirm that a part of the antiproliferative activity of QN-302 in a model of PDAC (MIA PaCa-2 cells) relies on its DNA damaging property. Then, we show that this activity could be potentiated when used in combination with an inhibitor of DNA repair, here the poly(ADP-ribose) polymerase-1 (PARP1) inhibitor Olaparib (*a*.*k*.*a*. AZD2281, or Lynparza, Fig. 1A) (35-37): this strategy, referred to as chemically induced synthetic lethality (CISL) (38, 39), greatly enhances the therapeutic potential of QN-302 and thus, opens new clinically-oriented possibilities.

## Results

### QN-302 is highly toxic in cultured cells

The antiproliferative activity of both QN-302 and PDS was first assessed in HeLa cells, used as a model for cancer cells (cervical cancer). Both ligands are quite active, with IC_50_ = 0.6±0.5 μM for PDS and 4.2±1.3 nM for QN-302, established using the sulforhodamine B (SRB) assay (40) after 72-h treatment (Fig. 1C, Additional file 1: Figure S1, Additional file 2: Table S1). The very potent antiproliferative activity of QN-302 was confirmed in 2 other cancer cell lines, with IC_50_ = 3.3±0.7 nM in MIA PaCa-2 cells (pancreatic cancer, further discussed below) and 4.6±1.2 nM in MCF-7 cells (breast cancer) and in the immortalized but non-cancerous HEK-293-T cells (human embryonic kidney), where it was found to be slightly less active (Additional file 1: Figure S1), with IC_50_ = 12.0±5.0 nM (*i*.*e*., a cancer *versus* non-cancer IC_50_ ratio between 3 and 9.5).

### QN-302 and PDS modulate G4 landscapes in HeLa cells

We first assessed if—and to what extent—both QN-302 and PDS target G4s in HeLa cells. The G4 engagement by PDS has already been established by optical imaging *via* different techniques, including *in situ* click imaging using the clickable PDS-α in both MRC5 (human fetal lung fibroblast) and U2OS cells (osteosarcoma) (29) or pre-targeted G4 imaging and/or *in situ* click imaging using labelable, biomimetic TASQ ligands (41) in both MCF-7 and HeLa cells (42, 43). We used here the latter method with the clickable ^az^MultiTASQ reagent (44). Cells were treated for a short incubation time (4 h) with ligands at a dose of *ca*. 5-fold IC_50_ (determined after 72-h treatments), which corresponds to a subtoxic concentration of QN-302 (20 nM) or PDS (2 μM) at this time point, for maximizing the sought-after outcomes while reducing unwanted secondary effects. Cells were then fixed (paraformaldehyde, PFA), blocked and incubated with ^az^MultiTASQ (1 h), which is then conjugated to the AlexaFluor-594 (AF-594) dye by copper-free click chemistry (strain-promoted alkyne-azide cycloaddition, SPAAC; 0.5 h) (45). Ascribing TASQ *foci* to *bona fide* G4 *foci* (see Methods and Additional file 1: Figures S2 and S3 for details about this quantification), we found that both ligands induce a global increase in G4 sites (Fig. 2). Because the effects monitored were subtle, experiments were performed as quadruplicates (>200 cells/experiment, 23 to 50 cells/conditions; see Additional file 3: Table S2) and two series of statistical analyses were performed, one for each experiment separately (Additional file 3: Table S2), and one on the pooled data (Fig. 2). On this basis, the number of G4 sites was found to increase upon treatment (*ca*. 3.8-fold with PDS, 4.5-fold with QN-302) with a difference in cellular distribution (nucleus *versus* cytoplasm).

**Figure 2:**
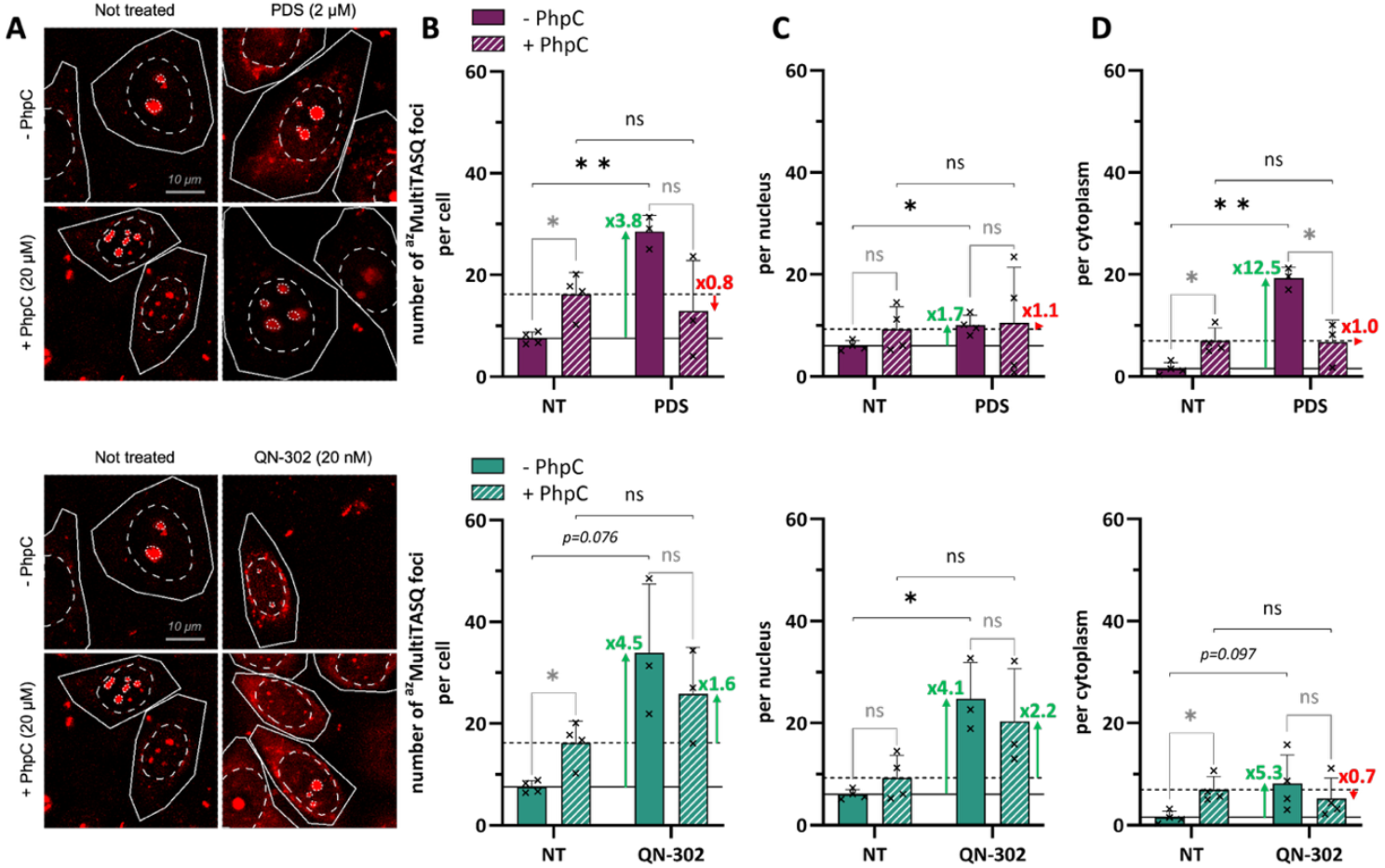
**A**. Representative images corresponding to the fluorescence detection of G4 *foci* (^az^MultiTASQ *foci*) in HeLa cells upon treatment with PDS (2 µM; upper panels) or QN-302 (20 nM; lower panels), with or without PhpC pre-incubation (20 µM). Images are maximal z-projection of 5 z-stacks (0.3 µm/z-stack) with total cell, nucleus and nucleolus outlined with respectively solid, dashed and dotted white lines, scale bar: 10 µm. **B**-**D**. Related quantification (either whole-cell (**B**), in the nucleus (**C**) or in the cytoplasm (**D**)) in HeLa cells (size selection: nuclear *foci* 3-100 voxels; cytoplasmic *foci* 20-100 voxels), untreated or treated with PDS (2 μM; upper panels) or QN-302 (20 nM; lower panels), without or with pre-incubation with PhpC (20 μM). Mean values and error-bars (standard deviation, sd) are represented as results collected from at least 3 independent experiments shown by individual dot (>200 cells/experiments, 20 to 50 cells/condition). Plain and dashed lines indicate the number of G4 *foci* in NT conditions without or with PhpC, respectively. Multiple unpaired t-test was employed for the statistical analyses with *: *P* ≤ 0.05, **: *P* ≤ 0.01, “ns” for non-significant: *P* > 0.05.

To further demonstrate G4 engagement by the ligands, we used PhpC, a prototype of a molecular helicase (34). Its ability to modulate the cellular G4 landscape downward was demonstrated by different techniques: optical imaging (with N-TASQ (46) in MCF-7 cells (47), BG4 (48) in HeLa cells (49) and QUMA-1 (50) in A549 cells) (51), G4RP-RT-qPCR (52, 53) (in both MCF-7 (47) and HT29 cells (colorectal cancer) (54)) and both RNA-seq and proteomics in patient-derived astrocytes (55). Here, we pre-treated HeLa cells with PhpC (20 μM, 1 h; Additional file 1: Fig. S1) before incubating them with QN-302 or PDS (20 nM and 2 μM respectively, for 4 h), and the cells were processed as above. The results seen in Fig. 2 (and Additional file 3: Table S2) show both a weak (*ca*. 1.5-fold in the nucleus) and unexpectedly strong increase (*ca*. 4.5-fold in the cytoplasm) in G4 sites upon PhpC treatment: this might originate in the ability of PhpC to foster several G4-involving biological pathways, notably cell stress response and RNA translation-related processes (documented in both astrocytes (55) and embryonic kidney cells, HEK-293-FT) (56). When co-incubated with G4 ligands, both PDS and QN-302 failed in raising significantly the number of *bona fide* G4 *foci*: as above, a quantification made on the basis of quadruplicates and a double statistical analysis (Additional file 3: Table S2) showed that none of the G4 ligand-triggered increase (from 0.7-to 2.2-fold) was statistically relevant, indicating that PhpC pre-incubation counteracts ligand-promoted G4 folding efficiently, thus lending strong support to the actual G4 nature of these *foci*.

Collectively, these results show that both PDS (confirmation) and QN-302 (demonstration) do engage G4s in cells, through an approach that abides by the very definition of chemical biology (57), in which chemicals are used as positive/negative modulators of biological systems to uncover cell circuitries where their intended targets (here, G4s) are involved (58, 59). These results highlight the strong effect of QN-302 on G4s, and more particularly DNA G4s, which led us to further explore the functional consequences of this interaction notably with regards to genomic instability.

### QN-302 and PDS trigger G4-mediated DNA damage in HeLa cells

The functional consequence of this interaction was further studied, notably to assess whether ligand-bound G4s compromise genomic stability. This has been already reported for PDS, whose DNA-damaging properties (*i*.*e*., its ability to trigger DSBs) are well-documented in a series of cell lines including U2OS (osteosarcoma), MDA-MB-231 (breast cancer) and HeLa cells (29, 32, 33, 43), and this was recently demonstrated for QN-302 in U2OS cells (60), with a particular focus on telomeres. Here, we confirmed this in HeLa cells, treating them with subtoxic concentrations (*vide supra*) of QN-302 (10, 20 and 50 nM) or PDS (2, 4 and 10 μM) for a short time (4 h) before fixation (PFA), blocking and incubation with antibodies raised against the phosphorylated H2AX histone (on Ser139) γH2AX, used as a marker of DNA breakage (61). An automatized quantification of γH2AX *foci* made on the basis of sextuplicates, and a double statistical analysis for some of them (>1000 cells/experiment, 76 to 848 cells/conditions; see Additional file 4: Table S4 and Fig. 3), indicate that both ligands induce strong increases of DSBs by *ca*. 1.7- and 7.2-fold with 4 and 10 μM PDS, respectively (no modifications with 2 μM PDS), and *ca*. 4.6- and 5.3-fold with 10 and 20 nM QN-302, respectively (a weak modification with 50 nM QN-302, which originates in the very low amount of surviving cells in these conditions). As above, we further demonstrated the G4 involvement in ligand-induced genetic instability by pre-treating HeLa cells with PhpC (20 μM, 1 h; −0.9±2.3 γH2AX *foci*/nucleus when used alone) before incubating them with QN-302 or PDS (4 h), and processing them as above. The results seen in Fig. 3 (and Additional file 4: Table S3) offer a clear evidence that PhpC efficiently mitigates G4-mediated genetic instability: when co-incubated with G4 ligands, both PDS and QN-302 raise the number of γH2AX *foci* in a modest and non-significant manner (from 0.3-to 3.7-(PDS) and 1.7-to 2.3-fold increase (QN-302), none of them being statistically significant). These results demonstrate that PhpC could prevent to a very promising extent the DNA damage mediated by the stabilization of DNA G4s by these ligands, highlighting its unique genoprotective properties.

**Figure 3:**
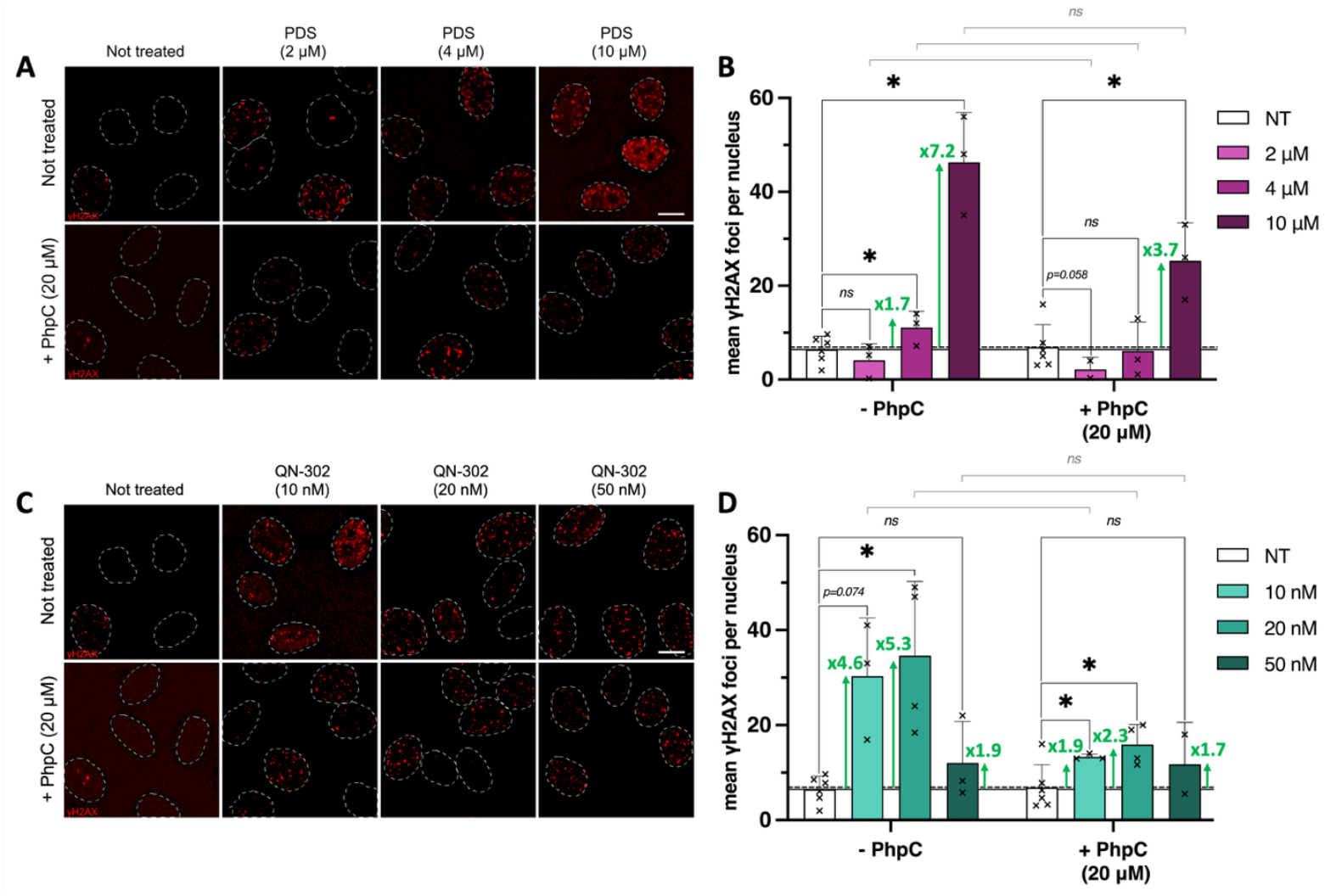
Representative images (**A, C**) and related quantification (**B, D**) corresponding to DNA damage (γH2AX) immunodetection experiments performed in HeLa cells (quantified as γH2AX *foci*/nucleus), either untreated or upon incubation with increasing concentrations of PDS (2 to 10 μM; upper panels) or QN-302 (10 to 50 nM; lower panels) without or with pre-incubation with PhpC (20 μM). Representative images are maximal z-projection of seven z-stacks (0.3 µm/z-stack) with nuclei outlined with dashed white lines, scale bar: 10 µm. Mean values of γH2AX *foci* and error-bars (standard deviation, sd) are represented as results collected from at least 3 independent experiments shown by individual dot. Plain and dashed lines indicate the number of γH2AX *foci* in NT conditions without or with PhpC, respectively. Multiple unpaired t-test was employed for the statistical analyses with *: *P* ≤ 0.05; “ns” for non-significant: *P* > 0.05.

### A competitive G4 binding study

We verified that the modulation of G4 landscapes described above does not originate in a simple competition of the G4-interacting compounds (QN-302 and PDS *versus* PhpC) for their targets. To this end, we established their affinity for G4s *in vitro* using two models: the c-Myc promoter (DNA G4) (62) and the NRAS 5’-UTR (RNA G4) (63). This affinity was established using two complementary *in vitro* assays: the fluorescence quenching assay (FQA) (64) and FRET-melting assay (65). The apparent G4 affinity (^app^*K*_D_) of QN-302 and PDS (Fig. 4A, Additional file 1: Fig. S4 and Additional file 5: Table S4) was found to be 0.21±0.03 µM (Cy5-Myc) and 1.65±0.15 µM (Cy5-NRAS) for QN-302 and 0.41±0.08 μM (Cy5-Myc) and 0.22±0.10 μM (Cy5-NRAS) for PDS, *versus* ≥ 100 μM for PhpC in both instances (47). Of note, the FQA experiments were performed in a reverse manner with QN-302, owing to its intrinsic fluorescence (see Methods). By FRET-melting experiments (Fig. 4B, Additional file 1: Fig. S5 and Additional file 5: Table S4), the thermal stabilization (ΔT_1/2_, using F-Myc-T) imparted by QN-302 and PDS (5 mol. equiv., 1 μM) was found to be 14.4±0.1°C and 28.7±0.8 °C for QN-302 and PDS, respectively, *versus* 0.2±0.1 °C for PhpC.

**Figure 4:**
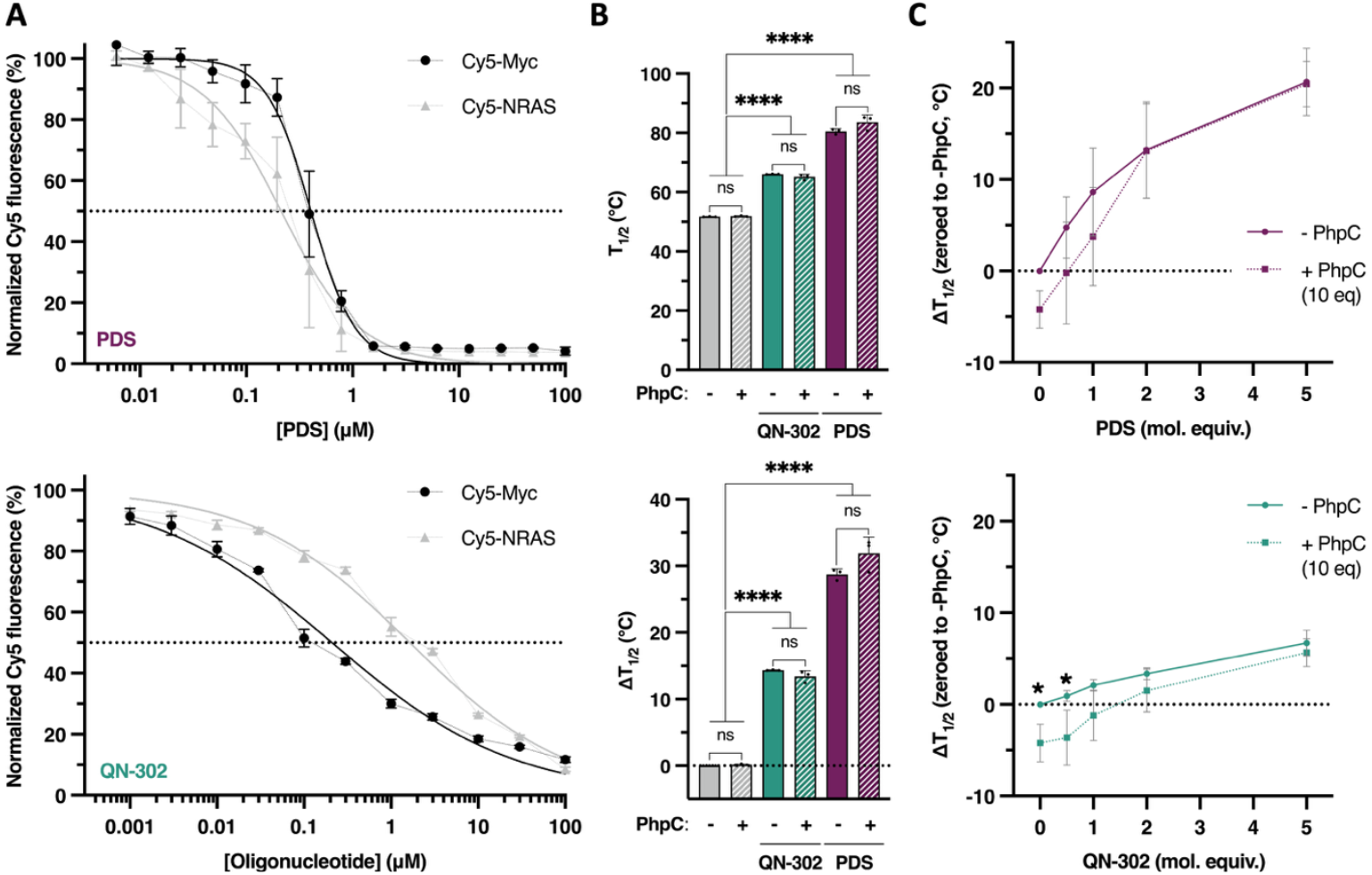
Quantification of the G4-interacting properties of PDS (upper panels) and QN-302 (lower panels) by either FQA (**A**) and competitive FRET-melting assays (**B, C**). FQA (for PDS) was performed with fixed oligonucleotide concentration (200 nM), being either DNA G4 (Cy5-Myc) or RNA G4 (Cy5-NRAS), and increasing concentrations of PDS (from 0.006 to 100 µM); reversed FQA (for QN-302) was performed with fixed ligand concentration (200 nM) and increasing concentrations of labeled oligonucleotides (from 0.001 to 100 µM), being either DNA G4 (Cy5-Myc) or RNA G4 (Cy5-NRAS). Two competitive FRET-melting experiments were performed: a classical competition (**B**) in which 0.2 μM DNA G4 (F-Myc-T) was incubated with 5 mol. equiv. of ligand (QN-302 or PDS, 1 µM), without or with 10 μM PhpC; a non-classical competition (**C**) in which 0.2 μM DNA G4 (F-Myc-T) was first pre-incubated overnight with 2 μM PhpC before being added of 0.5 to 5 mol. equiv. of ligand (QN-302 or PDS, 0.1 to 1 µM). The results are collected from triplicates (n = 3) across two to three independent studies. Two-way ANOVA was employed for the statistical analyses *: *P* ≤ 0.05, **: *P* ≤ 0.01, ***: *P* ≤ 0.001, ****: *P* ≤ 0.0001; “ns” for non-significant: *P* > 0.05.

This series of data showed that the G4 affinity of QN-302 and PDS is far higher than that of PhpC, making it unable to simply compete for G4 binding against established ligands. This was further demonstrated *via* competitive FRET-melting experiments in which the experiments were repeated in the presence of 2 μM PhpC (10 mol. equiv.): as seen in Fig. 4B (Additional file 1: Fig. S6 and Additional file 5: Table S4), the presence of PhpC slightly affected the ligand-triggered stabilization (ΔΔT_1/2_ = −0.94 °C for QN-302 and 3.13 °C for PDS). However, to demonstrate the ability of PhpC to distort G4s and thus, impede the proper interaction of QN-302 and PDS, we worked in a sequential manner (which mimics cell-based investigations), pre-folding a G4-forming sequence in the presence of PhpC (10 mol. equiv.) before performing a FRET-melting experiment (using F-Myc-T) with increasing concentrations of ligands (from 0 to 5 mol. equiv.). As seen in Fig. 4C (and Additional file 5: Table S4), the stability of the G4 itself was affected (ΔΔT_1/2_ = −4.20 °C), and the difference in ligand-triggered stabilization is maximal at low ligand concentrations (ΔΔT_1/2_ = −3.62 and −1.2 °C for 0.5 and 1.0 mol. equiv. QN-302; −0.23 °C for 0.5 mol. equiv. PDS), while being zeroed at high concentration (because of the difference in G4 affinity of QN-302 and PDS *versus* PhpC, *vide supra*). These results thus show that the modulation of the G4 landscape by pre-incubation with PhpC, as it is performed in cells, could be recapitulated *in vitro* provided that it is performed in a stepwise manner.

### The DNA damaging property of QN-302 is potentiated by Olaparib

The *in vitro* experiments described above have been performed in HeLa cells because we wanted to compare the properties of QN-302 *versus* PDS, which was extensively studied in this cell line (33, 43). To move closer to clinical trials, we next investigated the antiproliferative effects of QN-302 in PDAC cells (MIA-PaCa-2 cells) because this compound is currently in phase 1 clinical trial on PDAC patients (NCT06086522). More specifically, we wanted to assess whether QN-302 damages DNA in MIA PaCa-2 cells and the effect of a combination with Olaparib, used to inhibit DNA repair and thus, to potentiate the effects of QN-302. Olaparib is a PARP1 inhibitor, frequently used to induce synthetic lethality in homologous recombination (HR)-deficient cells. Olaparib, which delays single-strand DNA break (SSB) repair and generates DSBs, is FDA approved since 2014 (to treat ovarian cancer) (66, 67). It is often used in combination with DNA damaging agents to enhance their antiproliferative activity (68), notably to treat advanced ovarian and breast cancers as well as maintenance therapy in PDAC (69, 70). The interplay between G4s and PARP1 on one side, and G4 ligands and PARP1 inhibitors on the other side is complex, for several reasons (71): *i*-G4 ligands could be PARP1 inhibitors, and *vice versa* (see below); *ii*-a G4-forming sequence is found in the promoter region of PARP1, which might govern its expression (72); and *iii*-PARP1 itself could bind to G4-forming sequences (73, 74). From a small molecule point of view, a G4-interacting gold(I) complex was for instance shown to inhibit PARP1 activity *in vitro* (75), and a series of thioquinoline were developed to exert both G4-binding and PARP1-inhibiting activities (76). Conversely, PARP1 inhibitors such as Niraparib—but not Olaparib—were shown to interact with G4-forming sequences (77, 78). When both ligands and inhibitors are used in combination, synergistic effects have been monitored: this was pioneered with RHPS4 in combination with GPI-21016 in immortalized fibroblasts (BJ-HELT) and colorectal cancer (HT29 and HCT116 cells) (79), and confirmed more recently *via* a combination of CX-5461 and Talazoparib in ovarian cancer (OVCAR-8 cells) (80). Olaparib was previously used in combination with Ru complexes interacting with different DNA structures (including G4s) in breast cancer (MDA-MB-231 cells) (81).

Here, we thus verified whether the truly G4-selective QN-302 could act synergistically with Olaparib in PDAC (Fig. 5, Additional file 2: Table S1). This was done by detecting γH2AX *foci* in MIA PaCa-2 cells treated with toxic concentrations of QN-302 (5.0 nM, IC_50_ = 3.3±0.7 nM, Fig. 1) and Olaparib (50 μM, IC_50_ = 51.9±10.4 μM, Additional file 1: Fig. S1) for 24 h. When used alone, QN-302 does not induce DSBs (+2.5±4.0 γH2AX *foci*/nucleus) while Olaparib damaged DNA (+22.0±8.2 γH2AX *foci*/nucleus). When combined (Fig. 5A and Additional file 6: Table S5), the number of γH2AX *foci*/nucleus increased (+52.0±25.1 γH2AX *foci*/nucleus), which is *ca*. 2-fold beyond a simple additive effect, and thus, originates in synergy between the two drugs’ activity. This synergy was confirmed in HeLa cells, using the same concentration range (Fig. 5B and Additional file 6: Table S5). When used alone at this concentration, QN-302 did not trigger DSBs while Olaparib did (+2.3±3.8 and 13.0±3.0 γH2AX *foci*/nucleus, respectively) and the combination led to an increase in DSBs (+49.8±34.7 γH2AX *foci*/nucleus), which is *ca*. 3-fold beyond the additive effect.

**Figure 5:**
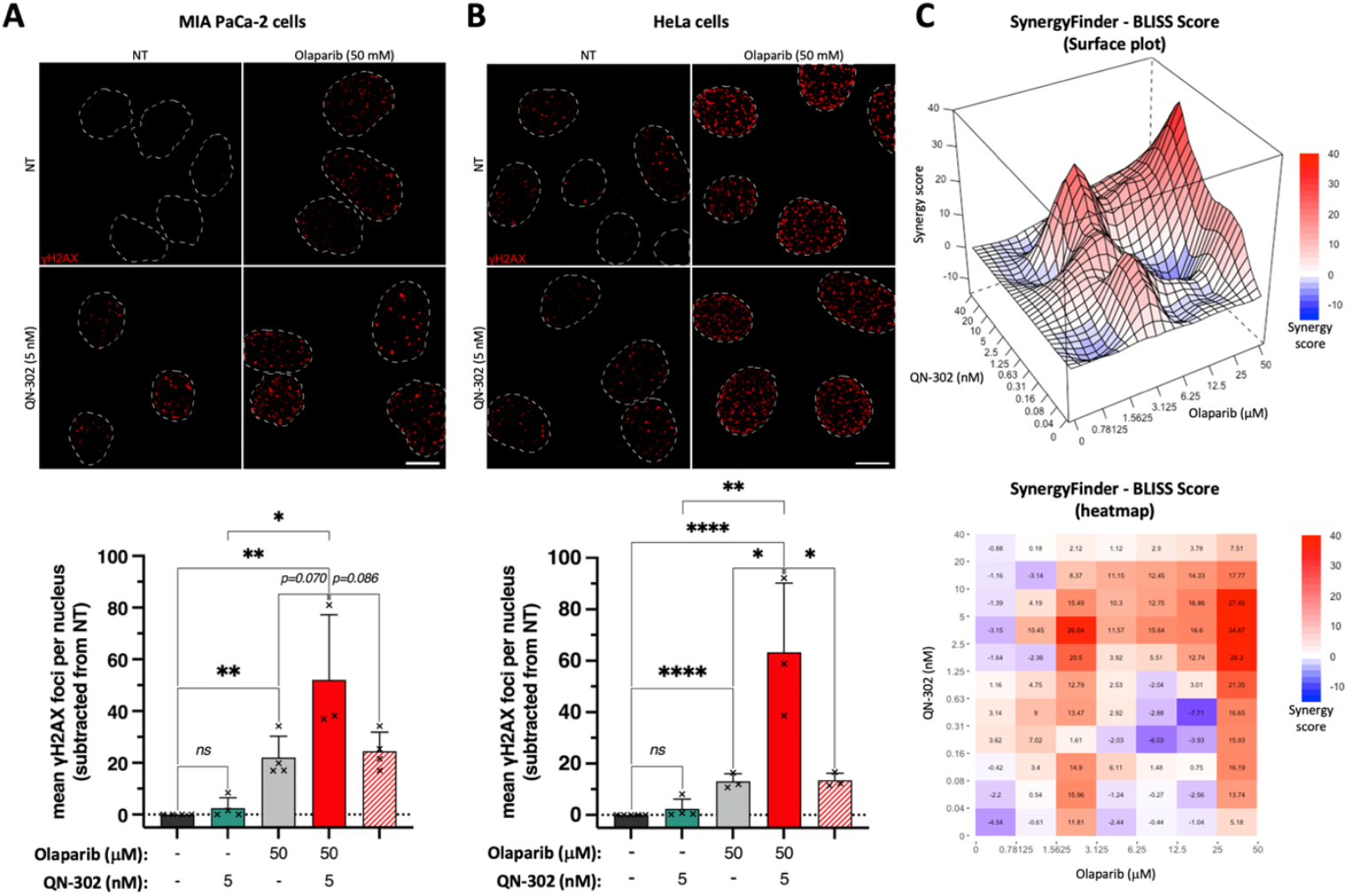
Study of the synergistic interaction between QN-302 and Olaparib in either MIA PaCa-2 (**A, C**) or in HeLa cells (**B**). **A-B.**Representative images (upper panels; maximal z-projection of 7 z-stacks (0.3 µm/z-stack) with nuclei outlined with dashed white lines, scale bar: 10 µM) and related quantification (lower panels; quantified as γH2AX *foci*/nucleus) of the synergy assessed by γH2AX immunodetection using 50 μM Olaparib and 5 nM QN-302. Mean values of γH2AX *foci*/nucleus (corrected to NT condition, indicated by dotted line) and error-bars (standard deviation, sd) are represented as results collected from at least 3 independent experiments shown by individual dot; red-dashed-bar represents the calculated theoretical additive effect. Fisher’s LSD tests were employed for the statistical analyses with *: *P* ≤ 0.05, **: *P* ≤ 0.01, ***: *P* ≤ 0.001; ****: *P* ≤ 0.0001; “ns” for non-significant: *P* > 0.05. **C**. Drug combination approach (assessed by SRB assay) in MIA PaCa-2 cells (from 0.04 to 40 nM of QN-302; from 0.78 to 50 μM of Olaparib). Mean values of BLISS score (calculated by SynergyFinder) are results from 3 independent experiments.

To confirm this synergy, we evaluated the toxicity (SRB assay, 72 h-treatment) of combinations of QN-302 (from 0.04 to 40 nM) and Olaparib (from 0.78 to 50 μM) in MIA PaCa-2 cells: both the 3-D plot and the heatmap seen in Fig. 5C (and Additional file 1: Fig. S7), generated using SynergyFinder Plus (82), illustrates the synergistic (red) or antagonistic (blue) interactions between the antiproliferative effects of two compounds. It is accepted that the synergistic combinations are those with a BLISS Score >10; here, the best combinations are obtained with 5 – 20 nM QN-302 and 12.5 – 50 μM Olaparib (BLISS Score > 12.4). A quite interesting combination (BLISS Score = 26.0) is obtained with 5 nM of QN-302 and 3.12 μM Olaparib, which deserves to be further investigated, with an eye towards clinical applications.

To correlate these observations with ligand-associated genetic instability, we verified the induction of γH2AX *foci* by the four possible combinations of high/low concentrations of QN-302/Olaparib after 24 h-treatment (Fig. 5, Additional file 1: Fig. S8 and Additional file 6: Table S5): for instance, the synergistic combinations [5 nM QN-302+50 μM Olaparib] (BLISS Score = 34.9) triggers a notable increases in DSBs (+52.0±25.1 γH2AX *foci*/nucleus, respectively), while the antagonistic combination [0.04 nM QN-302+0.78 μM Olaparib] (BLISS Score = −4.5) triggers a modest increase only (+4.3±2.9 γH2AX *foci*/nucleus, respectively).

Finally, we tried to confirm these results in HeLa cells, which was possible for the immunodetection of DNA damage (Figure 5B): for instance, using the same concentration ranges, the synergistic combinations [5 nM QN-302+50 μM Olaparib] and [0.04 nM QN-302+50 μM Olaparib] led to a notable increase in DSBs (+51.5±35.5 and 21.5±15.9 γH2AX *foci*/nucleus, respectively), while other combinations ([5 nM QN-302+0.78 μM Olaparib] and [0.04 nM QN-302+0.78 μM Olaparib]) do not (+6.7±9.6 and 5.6±5.8 γH2AX *foci*/nucleus, respectively). However, this synergy was poorly confirmed by toxicity studies (Additional file 1: Figs. S9 and S10 and Additional file 6: Table S5): even if some local maxima were obtained (*e*.*g*., BLISS Score = 11.0 with [0.16 nM QN-302+12.5 μM Olaparib] or 13.7 with [0.31 nM QN-302+1.56 μM Olaparib]), the mean score over the matrix is very low (0.7 *versus* 6.6 for MIA-PaCa-2 cells), with a poor statistical relevance (*p* = 3.31e-01 *versus* 7.66e-9 for MIA-PaCa-2 cells). The origin of this lack of synergy remains to be deciphered, but might be two-fold: first, a difference in both DNA damage response and cell cycle regulation between the two cell lines, related to their retinoblastoma protein (pRb) status, HeLa cells being pRb-deficient cells and MIA-PaCa-2 cells pRb-proficient (83, 84). pRb is a tumor suppressor gene (85); its inactivation impairs both DNA damage repair and cell cycle control (86, 87). Our hypothesis is that the pRb status makes HeLa cells less sensitive to the QN-302/Olaparib combination than MIA-PaCa-2 cells. Second, these results must be seen in the context of cell doubling times (16 *versus* 30 h for HeLa and MIA PaCa-2 cells, respectively): this implies 4.5 HeLa *versus* 2.4 MIA PaCa-2 cell doublings during the experiments (72 h), meaning that surviving cells learned how to deal with ligand-induced genetic instability.

## Conclusion

QN-302 results from the structural optimization of naphthalene diimide (NDI)-based ligands, a popular scaffold for targeting G4s (88). Specifically, QN-302 is a four-armed NDI which was found to be the most active against PDAC cell lines (MIA PaCa-2, PANC-1, CAPAN-1 and Bx-PC3, with IC_50_ in the low nM range) (22, 24). Its favorable pharmacokinetics characteristics (*e*.*g*., a half-life of 37 h *in vivo*) made it suited to be evaluated in several small-animal models of PDAC, where it displays significant anti-tumor activity (for example against xenografted mice) at low doses (1 mg/mL, once weekly over 4 weeks). QN-302 was granted Orphan Drug designation by the FDA in Dec. 2022 and investigational new drug (IND) clearance in Aug. 2023 by the FDA for phase I clinical trials (89). These investigations, currently ongoing, are based on 28-day once-weekly dose escalation; when used at the lowest dose, treated patients showed no significant adverse effects.

To go a step further in this clinical exploitation, combination drug therapy is an appealing way to use lower individual drug doses as a result of a possible synergy during which drugs used in combination enhance the efficacy of each other (thus minimizing side effects). The greatest challenge is to find good drug partners, among hundreds of possible combinations: we have identified here the DNA damaging properties of QN-302 as a possible cellular pathway to be exploited for combination therapy. We first demonstrated that DNA breaks induced by QN-302 do involve G4s, due to a quite unique chemical biology strategy based on the molecular helicase PhpC. Then, we associated QN-302 with a clinically approved inhibitor of DNA repair, Olaparib, with the aim of synergizing its antiproliferative activity: this strategy was validated *in vitro*, studying MIA PaCa-2 cells in culture in multi-well plates allowing for simultaneous measurement of the effect of many drug combination ratios. We found a very interesting synergy between QN-302 and Olaparib, with >20 combinations displaying a BLISS Score >10 (and 6 combinations with a score >20), thereby providing a solid basis from which clinical trials could be envisaged. These results thus provide fertile starting points for further developments aiming at potentiating the therapeutic use of QN-302 in the clinic for the treatment of PDAC and potentially other human cancers (90).

## Methods

### Reagents

QN-302 was solubilized at 5 mM in 25% DMSO, PDS at 20 mM in DMSO, PhpC at 20 mM in DNase/RNase-free ultra-pure water; ^az^MultiTASQ at 2 mM in DNase/RNase-free ultra-pure water. All oligonucleotides used in this study were purchased from Eurogentec (Belgium) (see Additional file 1), diluted in ultrapure water (18.2 MΩ.cm resistivity) at 500 µM for stock solutions and stored at −20 °C. The actual concentration of these stock solutions was determined through a dilution to 1 to 5 μM theoretical concentration *via* a UV spectral analysis at 260 nm with the molar extinction coefficient values provided by the manufacturer.

### Cell lines and cell culture

MCF-7, HeLa, MIA PaCa-2 and HEK-293-T cells were routinely cultured in 75 cm^2^ tissue culture flasks (Nunc) at 37 °C in a humidified 5% CO_2_ atmosphere in Dulbecco’s Modified Eagle Medium (DMEM, Dutscher, cat. n° L0104) supplemented with 10% (v/v) fetal bovine serum (FBS, Dutscher, cat. n° S1810) and 1% (v/v) Penicillin-Streptomycin (Pen-Strep, 106.8 U/mL Pen, 106.8 µg/mL Strep, Gibco, cat. n° 151-40-122).

### SRB assay

According to Ref. (40). Experiments were performed in 96-well plates (flat bottom; Corning-Falcon cat. n° 353072) using CLARIOstar Plus plate-reader device (BMG LABTECH). Cells were seeded in a 96 well-plate at a density of 3,000 (MCF-7 and HeLa) and 5,000 cells/well (MIA PaCa-2 and HEK-293-T) with a final volume of 100 µL, and allowed to recover 24 h. Cells were treated live with various concentration of ligands (100 µL of 2X ligands in supplemented DMEM) or not (negative control = untreated cells; only supplemented DMEM plus the same percentage of DMSO, if applicable), for 72 h at 37 °C. Cells were fixed with an addition of 100 µL of cold 10% (w/v) trichloroacetic acid (TCA) to wells for 45 min at 4 °C, washed with tap water (x3), allowed to dry overnight at 25 °C, stained with 0.057% (w/v) SRB (in 1% (v/v) acetic acid) for 30 min at 25 °C under agitation, then washed with 1% (v/v) acetic acid (x3) and allowed to dry overnight at 25 °C. After incubating the plate with cold 10 mM unbuffered Trizma base solution for 30 min at 25 °C under agitation, absorbance (λ= 490-540 nm; with 10 nm step) was recorded. Final data were analyzed with Excel (Microsoft Corp.) and GraphPad Prism version 9.5.1 for Mac OS (GraphPad software). Mean absorbance values, normalized (to NT condition) % cell viability and then IC_50_ values were calculated on the basis of at least 3 independent experiments.

### Optical imaging

MIA PaCa-2 and HeLa cells were seeded on coverslips in 24-well plates at a concentration of 50,000 cells/well and 100,000 cells/well, respectively. Cells were allowed to attach for 40 (MIA PaCa-2) or 24 h (HeLa). Then, cells were either pre-treated or not with PhpC (20 µM; 1h) and treated for 4 h with PDS or QN-302 with indicated concentrations; or cells were co-treated for 24 h with or without Olaparib and with or without QN-302 with indicated concentrations. Cells were washed 3 times with cold PBS and prepared for different immuno-or chemo-detection experiments. After cold PBS washing steps, pre-extraction was performed on ice twice for 3 min with ice-cold pre-extraction buffer (10 mM PIPES pH 7, 100 mM NaCl, 300 mM sucrose, 3 mM MgCl_2_) supplemented with 0.3% or 0.7% (v/v) of Triton X-100, respectively for MIA PaCa-2 or HeLa cell lines. After pre-extraction steps, cells were washed once with cold PBS and fixed, for 10 min at room temperature (RT) with PFA (28908, Themo Scientific) diluted in PBS (2% PFA for γH2AX immunodetection or 4% PFA for ^az^MultiTASQ chemodetection). For immunodetection of γH2AX: after fixation, cells were washed three times with PBS and permeabilized with PBS-Triton X-100 0.1% (v/v) for 10 min. Cells were washed three times with PBS and incubated with blocking buffer (PBS, 1% (w/v) BSA, 0.2% (w/v) fish gelatin, 0.1% (v/v) Triton X-100) for 30 min at room temperature prior to a 2-h incubation at RT with appropriate dilution (1/2000) in blocking buffer of primary antibody (anti-phospho-histone H2A.X (Ser139) mouse mAb clone JBW301, 05-636, Merck Millipore (Sigma)). Cells were washed five times with TBS-Tween20 0.1% (v/v) then incubated with appropriate AlexaFluor(AF)-coupled secondary antibody (anti-mouse IgG (H+L) (Alexa Fluor 594 conjugated), A11032, Invitrogen) diluted at 1/2000 in blocking buffer for 45 min at room temperature. Cells were washed twice with TBS-Tween20 0.1% (v/v) and incubated with 1 µg/mL of DAPI (D9542-10MG, Sigma-Aldrich) in PBS-Triton X-100 0.1 % (v/v) for 15 min at room temperature. Coverslips were assembled on microscope glass slide with Vectashield^®^ (H-100-10, Vector Laboratories) and fixed with clear nail polish. According to Ref. (64), for chemodetection experiments: pre-extraction, cell fixation and permeabilization were performed as previously described (*vide supra*). Incubation with FBS-based blocking buffer (2% (v/v) FBS in PBS-Triton X-100 0.1 % (v/v)) was performed for 30 min at 37 °C for ^az^MultiTASQ staining. For G4 detection, 200 µL of 20 µM ^az^MultiTASQ solution (in FBS-based blocking buffer) was added to each well and incubated for 1 h at 37 °C. Cells were washed five times with TBS-Tween20 0.1% (v/v), before performing strain-promoted azide-alkyne cycloaddition (SPAAC): cells were incubated with “click” solution (0.5 µM DIBO-AF594 in FBS-based blocking buffer) for 30 min in the dark at room temperature. Finally, washing steps, DAPI staining, and coverslip mounting were performed as previously described for immunofluorescence staining. Epi-fluorescence acquisition of z-stacked images (0.3 µm/stack) was performed with Multiple Image Acquisition program of Olympus IX 83 widefield microscope equipped with 60X oil-objective and with excitation/emission filters for DAPI and TexasRed. After acquisition, images were deconvoluted with Weiner program (CellSens software, Olympus) and z-stack images were analyzed for quantification and representative images (z-stack = 5 or 7, maximum z-projection) were processed with FIJI software.

### ^*az*^MultiTASQ chemodetection quantification

^az^MultiTASQ quantification was performed as previously described (43). Since cytoplasmic staining was too faint in non-treated condition, manual segmentation of cells was implemented as a new user-friendly macro (see: https://github.com/ICMUB/azMultiTASQ_manually-outline). Briefly, after removal of ^az^MultiTASQ background noise (determined in region without any cells) cells were manually segmented (n= 50 cells/condition/experiment) and selected as regions of interest (ROI) for 5 consecutive z-stacks. Fluorescent signals (DAPI and TexasRed) were cleared outside the ROI (Figure S2) to avoid the quantification of multiple cells, and nucleus within ROI was automatically segmented using DAPI staining Figure S2). Then, ^az^MultiTASQ staining was automatically segmented (using the same parameters throughout all conditions from a given experiment) to obtain individual events of interest (EOI) considered as ^az^MultiTASQ *foci* (grey arrows, Figure S2). ^az^MultiTASQ segmented *foci* overlapping with segmented DAPI staining were separated from those within the cytoplasm, and ^az^MultiTASQ staining within nucleoli was automatically segmented and removed from the EOI within nucleus, as morphologically determined nucleoli correspond to nuclear ^az^MultiTASQ *foci* with a size > 100 voxels (Figure S2). Therefore, ^az^MultiTASQ nuclear *foci* quantified are small (3 to 100 voxels) and individual events specifically found in the nucleus (Figure S2). Segmented cytoplasmic ^az^MultiTASQ *foci* of interest were considered within a size from 20 to 100 voxels (Figure S2), as smaller *foci* could be due to background noise and larger *foci* could be hard to be separated. At the end of the quantification, both volume and number data of segmented events and of fluorescence intensity (within segmented events) were available for nucleolar, nuclear and cytoplasmic ^az^MultiTASQ staining, as well as those for DAPI staining. This set of data was used to determine outliers (*e*.*g*., DAPI staining missing or lowered, nucleoli not properly segmented, etc.) and to decipher variations between conditions (Figure S3). Of note, selected features for an automated segmentation of DAPI and of ^az^MultiTASQ staining were consistent throughout all conditions from a given experiment (see Figure S2), therefore overcoming variation of fluorescence intensity for quantified signals. Also, the number of ^az^MultiTASQ *foci* determined by this pipeline is not an absolute value and should be considered as a tool to reveal trends, as shown for one representative experiment in Figure S3. Quantification was performed on at least three independent experiments for each treatment and quantification.

### Fluorescence quenching assay

According to Ref. (64) for PDS. Calculation of the apparent dissociation constant ^APP^*K*_D_ is assessed through the ability of the ligand to bind to, and quench the fluorescence of a G4-forming sequence functionalized with a Cyanine 5 (Cy5) on its 5’ end. Each Cy5-labelled oligonucleotide (Eurogentec) was resuspended at 500 µM in ultrapure water and then diluted at 50 µM in CacoK10 buffer (10 mM lithium cacodylate pH 7.2, 10 mM KCl, 90 mM LiCl) in order to be folded. Cy5-labelled oligonucleotides were folded by heating at 90 °C for 5 min, then quickly cooled down on ice and kept at 4 °C the day before FQA experiment. Classical FQA (with PDS): experiments were performed in a 96-well plate (10762394, Greiner) with a final volume of 50 µL of Cy5-labelled G4 at a final concentration of 200 nM prepared in KD buffer (50 mM Tris-HCl pH 7.2, 150 mM KCl, 0.05 % Triton X-100). Serial dilutions of PDS ranging from 1000 to 0.06 µM (10X) were prepared in KD buffer and 5 µL were added in each well (1X final). Reverse FQA (with QN-302): experiments were performed in a 96-well plate with a final volume of 50 µL of QN-302 at a final concentration of 200 nM prepared in KD buffer. Serial dilutions of appropriate Cy5-oligonucleotide ranging from 1000 to 0.01 µM (10X) were prepared in KD buffer and 5 µL were added in each well (1X final). In both instances: fluorescence of Cy5 (λ_ex_: 595-625 nm; λ_em_: 650-700 nm) was measured with a CLARIOstar spectrophotometer before the addition of ligands (T0) and after 60-min incubation (T60) with gentle agitation in the dark. Mean values and standard deviation (SD) of at least 3 independent experiments performed in technical duplicates were plotted in function of ligand or DNA concentration. ^APP^*K*_D_ has been determined using a non-linear regression with a four-parameters dose-response curve in GraphPad Prism (v 10.1.2).

### FRET-melting assay

100 µM solutions of QN-302, PDS and PhpC were prepared in ddH_2_O. Experiments were performed in 96-well plates using Mx3005P qPCR device (Agilent) equipped with FAM filters (λ_ex_ = 492 nm; λ_em_ = 516 nm). Classical competition: to a mixture of F-Myc-T (final concentration: 200 nM) in CaCoK1 buffer (10 mM lithium cacodylate pH 7.2, 1 mM KCl, 99 mM LiCl) were added 5 mol. equiv. (1 µM) of either QN-302 or PDS, without or with 50 mol. equiv. (10 µM) PhpC. For the classical competitive FRET-melting protocol: after an initial increase step (from 25 to 90 °C, 30 s), a stepwise decrease (1 °C every 30 s for 67 cycles, from 90 to 25 °C) was performed followed by a stepwise increase (1 °C every 30 s for 67 cycles, from 25 to 90 °C), and measurements were made at the end of each cycle. Non-classical competition: a mixture of F-Myc-T (20 µM) was prefolded (5 min at 90 °C) in the presence of 10 mol. equiv. (200 µM) PhpC (or without, as control) in CaCoK 1 buffer and then cooled down overnight at 4°C. Prefolded F-Myc-T with or without PhpC was then diluted at 0.2 μM (final concentration of oligonucleotide) before the addition of 0.5, 1, 2, or 5 mol. equiv. (0.1, 0.2, 0.4, or 1 µM) of either QN-302 or PDS. Only the stepwise increase was performed (1 °C every 30 s for 67 cycles, from 25 to 90 °C), and measurements were made at the end of each cycle. For both instance: final data were analyzed with Excel (Microsoft Corp.) and GraphPad Prism (v 10.1.2). The emission of FAM was normalized (0 to 100%) and the T_1/2_ (°C) and ΔT_1/2_ (°C) were calculated (ΔT_1/2_ (°C) = (T_1/2 DNA + G4 ligand_) – (T_1/2 DNA alone_)) on the basis of three independent experiments performed in technical triplicates.

### Synthetic lethality studies

Synergistic assays were performed with simultaneous co-treatment of appropriate concentrations of QN-302 and Olaparib for 72 h either in MIA PaCa-2 or HeLa cells. Cell viability for each co-treatment was assessed by SRB staining, as previously described. Synergistic cell death (at least 3 independent experiments performed in technical replicates) was performed using SynergyFinder Plus and the SynergyFinder R package web application (https://www.bioconductor.org/packages/release/bioc/vignettes/synergyfinder/inst/doc/User_tutorual_of_the_SynergyFinder_plus.html#reshaping-and-pre-processing).

Synergistic cell death was also calculated as a percentage of additional lethality (see Figures S5 and S7) by subtracting each cell viability value to the untreated condition. Mean values of synergistic cell death (%) from at least three independent experiments (in technical duplicates) were plotted as a heatmap in GraphPad Prism (v 10.1.2).

## Supplementary information

**Additional file 1: Figure S1**: cytotoxicity curves obtained with QN-302, PDS and Olaparib; **Figure S2**: details about the quantification of ^az^MultiTASQ *foci* in HeLa cells treated by QN-302 or PDS, without or with pre-incubation of PhpC; **Figure S3**: representative example of quantification of ^az^MultiTASQ *foci* in HeLa cells treated by QN-302 or PDS, without or with pre-incubation of PhpC; **Figure S4**: FQA curves obtained with QN-302 and PDS; **Figures S5** and **S6**: FRET-melting curves obtained with QN-302 and PDS, without or with PhpC; **Figure S7**: synergistic interactions between QN-302 and Olaparib assessed by the SRB assay in MIA PaCa-2 cells; **Figure S8**: synergistic interactions between QN-302 and Olaparib assessed by the immunodetection of DSBs in MIA PaCa-2 cells; **Figure S9**: synergistic interactions between QN-302 and Olaparib assessed by the SRB assay in HeLa cells; **Figure S10**: synergistic interactions between QN-302 and Olaparib assessed by the immunodetection of DSBs in HeLa cells; **Additional file 2: Table S1**: determination of IC_50_ values by SRB; **Additional file 3: Table S2**: *in situ* click imaging for the quantification of G4 *foci*; **Additional file 4: Table S3**: immunodetection of ligand-induced DSBs; **Additional file 5: Table S4**: characterization of the G4-affinity of small molecules *in vitro*; **Additional file 6: Table S5**: synergistic interactions between QN-302 and Olaparib assessed by the immunodetection of DSBs in MIA PaCa-2 cells.

## Acknowledgment

The authors thank Marc Pirrotta (UBE Dijon, FR) for his help for the *in vitro* assays, Ibai Valverde and Sandy Raevens (UBE Dijon, FR) for the preparation of ^az^MultiTASQ, along with Pauline Lejault (UBE Dijon, FR) and Judy M. Y. Wong (UBC Vancouver, CA) for fruitful discussions.

## Authors contribution

D.M. and S.N. managed the project. G.P., A.P. and D.M. designed the experiments. G.P., A.P. and M.B. conducted the experiments. G.P., A.P. and D.M. processed the experimental data and analyzed the results. R.H.E.H. provided the consortium with PhpC, S.N. with QN-302. S.N. and D.M. wrote the manuscript.

## Funding

This work was supported by the Agence Nationale de la Recherche [InJUNCTION, ANR-22-CE44-0039-01] to D.M. for A.P. and by the Région BFC through the ICE PhD program [IMPALA; OPE-2024-0046-D103] to D.M. for G.P.

## Availability of data and materials

All data generated or analyzed during this study are included in this published article and its supplementary information files.

## Declarations Ethics approval

Not applicable.

## Consent for publication

All authors read and approved the final manuscript.

## Competing interests

S.N. is currently a member of the Scientific Advisory Board of Qualigen Therapeutics Inc. D.M. and the CNRS have licensed TASQs to Merck KGaA for commercialization; D.M. also provides consulting services for Idylle with the commercialization of PhpC.

---

Proposal for an image for TOC and/or display on webpages/social media

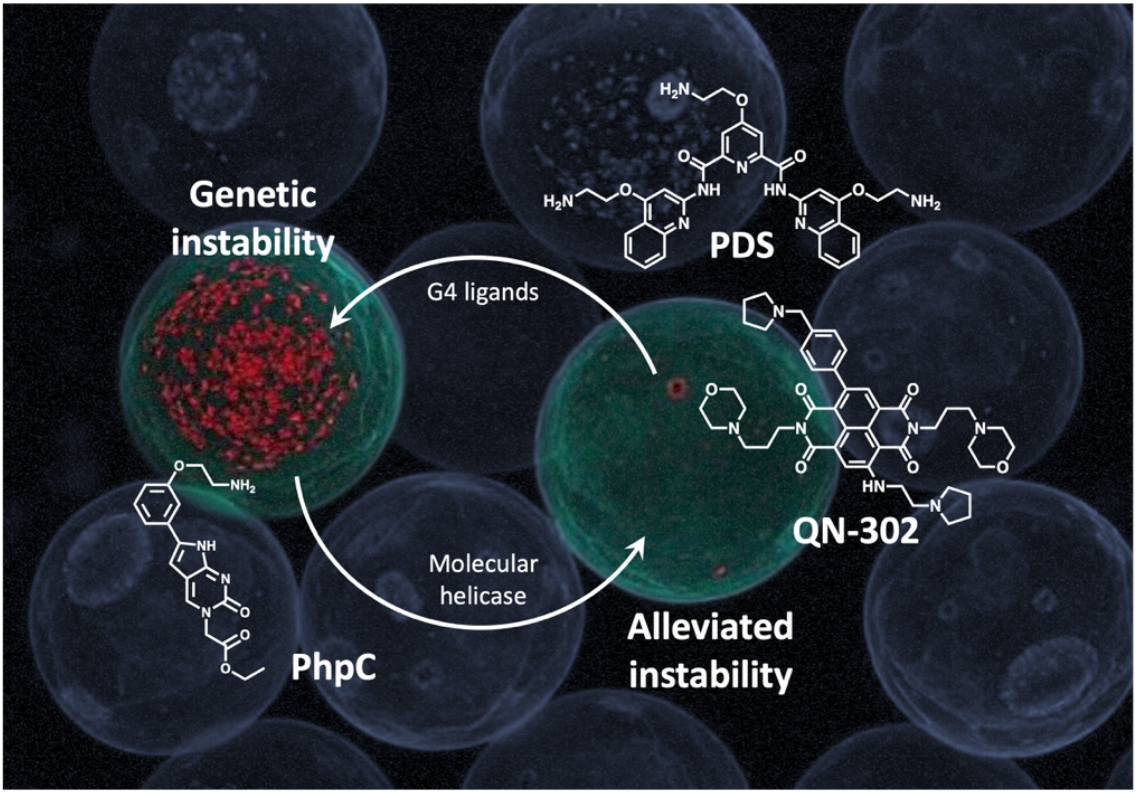

