## Additional file 1 for "The DNA damaging properties of the experimental G-quadruplex (G4) drug QN-302 are potentiated by the DNA repair inhibitor Olaparib and mitigated by the molecular helicase PhpC"

### Additional figures

Oligonucleotides used in this study:

| Oligonucleotide | Sequence |
|---|---|
| Cy5-Myc | Cy5-d[GAG-GGT-GGG-GAG-GGT-GGG-GAA-G] |
| Cy5-NRAS | Cy5-[GGG-AGG-GGC-GGG-UCU-GGG] |
| F-Myc-T | FAM-d[GAG-GGT-GGG-GAG-GGT-GGG-GAA-G]-TAMRA |

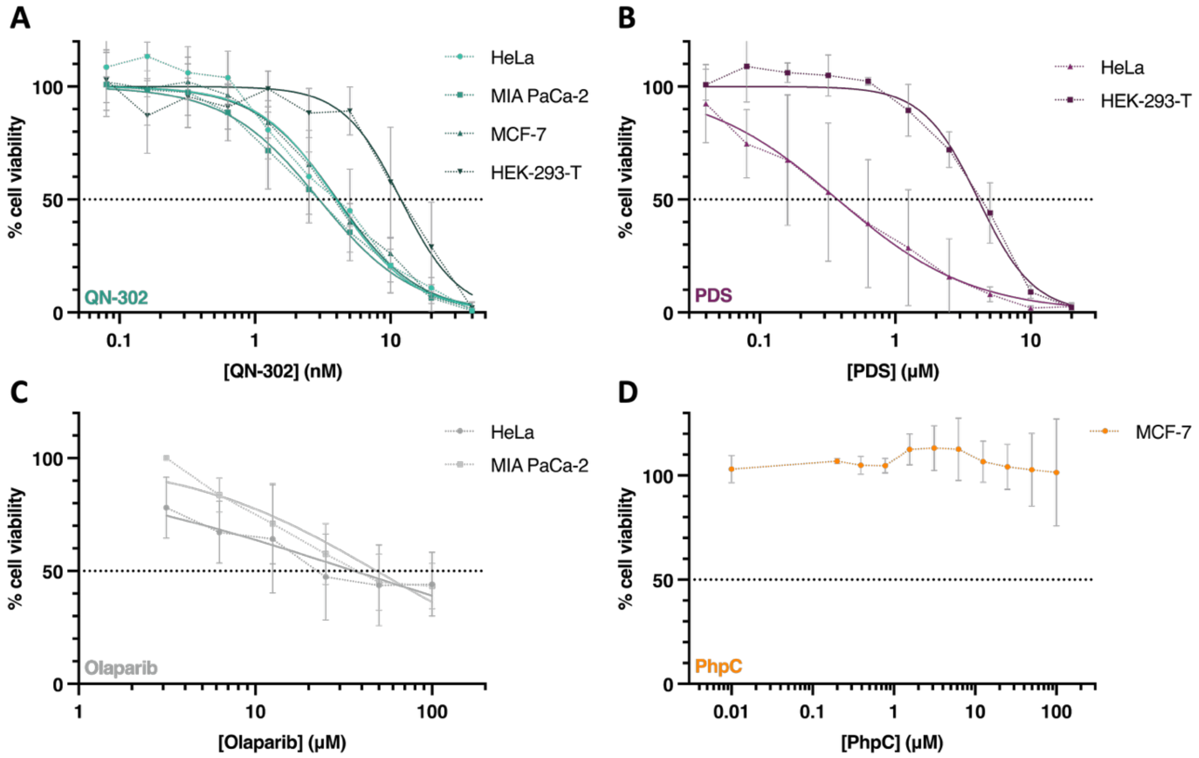


**Figure S1**: Cytotoxicity assays performed with immortalized (HEK-293-T cells) or cancer cell lines (HeLa, MCF-7 and MIA PaCa-2 cells) and PDS (up to 20 µM, **A**), QN-302 (up to 40 nM, **B**), Olaparib (up to 100 µM, **C**) and PhpC (up to 100 µM, **D**), established by the SRB assay after 72-h treatment. The results are collected from triplicates (n = 3) across three independent studies (n = 3).

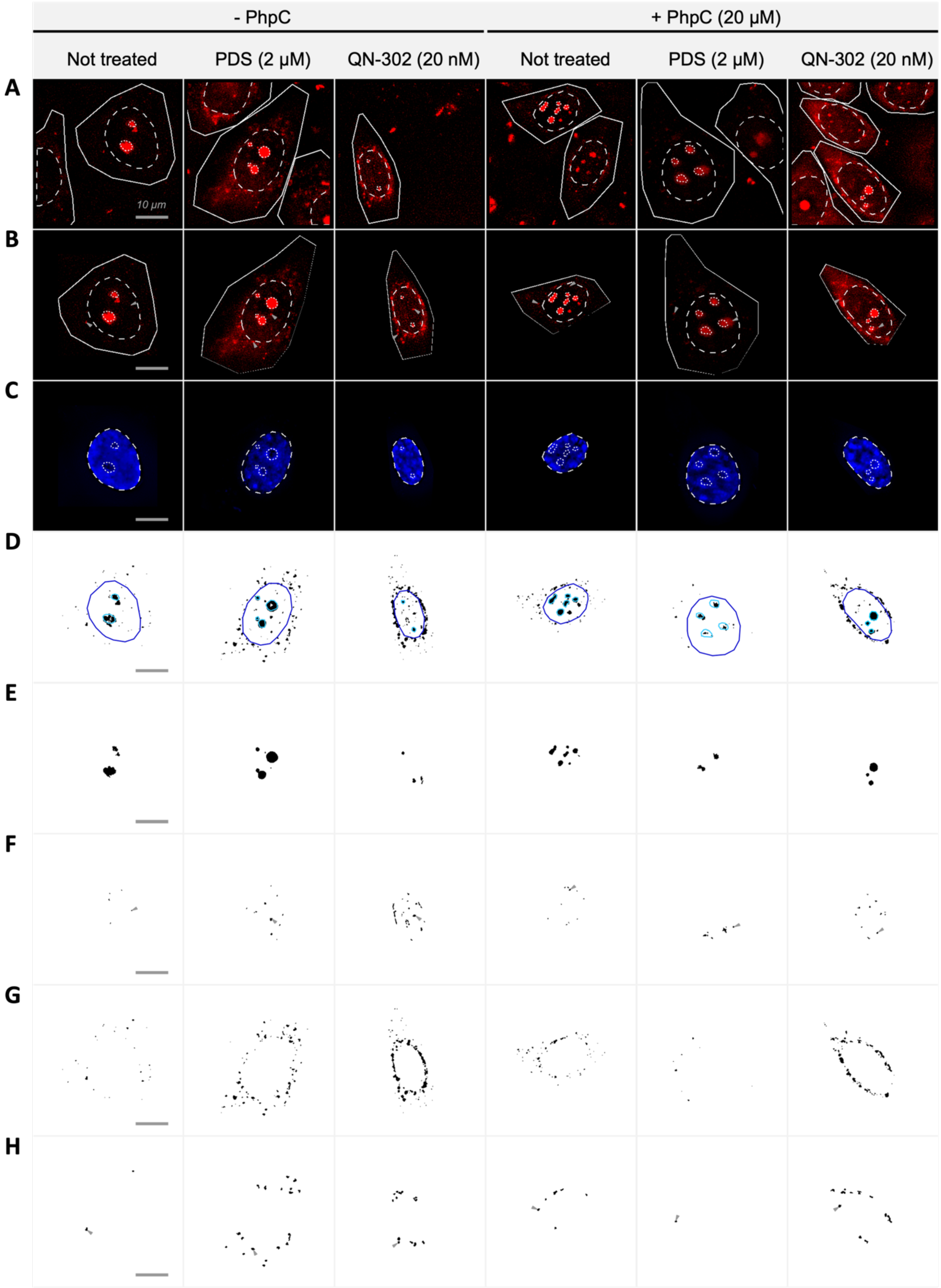


**Figure S2:** Representative images (used in **Fig. 2**) showing $^{az}$MultiTASQ quantification steps corresponding to each condition (with or without pre-incubation of PhpC (20 µM) and with or without G4-ligand treatment (PDS 2 µM or QN-302 20 nM)). Grey arrows (seen in **B, F** and **H**) show events of interest as example of objects to be quantified in nucleus (seen in **F**) and in cytoplasm (seen in **H**). Representative images are shown as maximal z-projection of 5 z-stacks (0.3 µm/stack) although quantification was performed from XYZ images. Scale bar corresponds to 10 µm. **A.** Representative images (after background noise removal) with white outlines showing *i)* cellular manual segmentation (considered as ROI) in solid line; *ii)* nuclear automated segmentation (from DAPI staining shown in **C**) in dashed line; and *iii)* morphologically determined nucleoli (overlapping with nuclear TASQ foci with size >100 voxels shown in **E**) in dotted line. **B.** ROI selection and clearing outside selected ROI to analyse only selected cell. **C.** DAPI staining allowing automated segmentation of nucleus outlined in white dashed line and morphological determination of nucleoli outlined in white dotted line. **D.** Automated segmentation of $^{az}$MultiTASQ foci (threshold value common to all conditions) with nucleus automated segmentation outlined in blue and with nucleoli morphological determination outlined in light blue. **E.** Automated segmentation of big (>100 voxels) nuclear TASQ foci perfectly fits with morphological determination of nucleoli. **F.** Selection (by overlapping with DAPI segmented nucleus) of small (3-100 voxels) nuclear TASQ foci. **G.** Selection of cytoplasmic TASQ foci (by overlapping with manual determination of ROI and removal of nucleus segmentation) without size-discrimination. **H.** Selection of medium (20-100 voxels) cytoplasmic TASQ foci to be quantified.

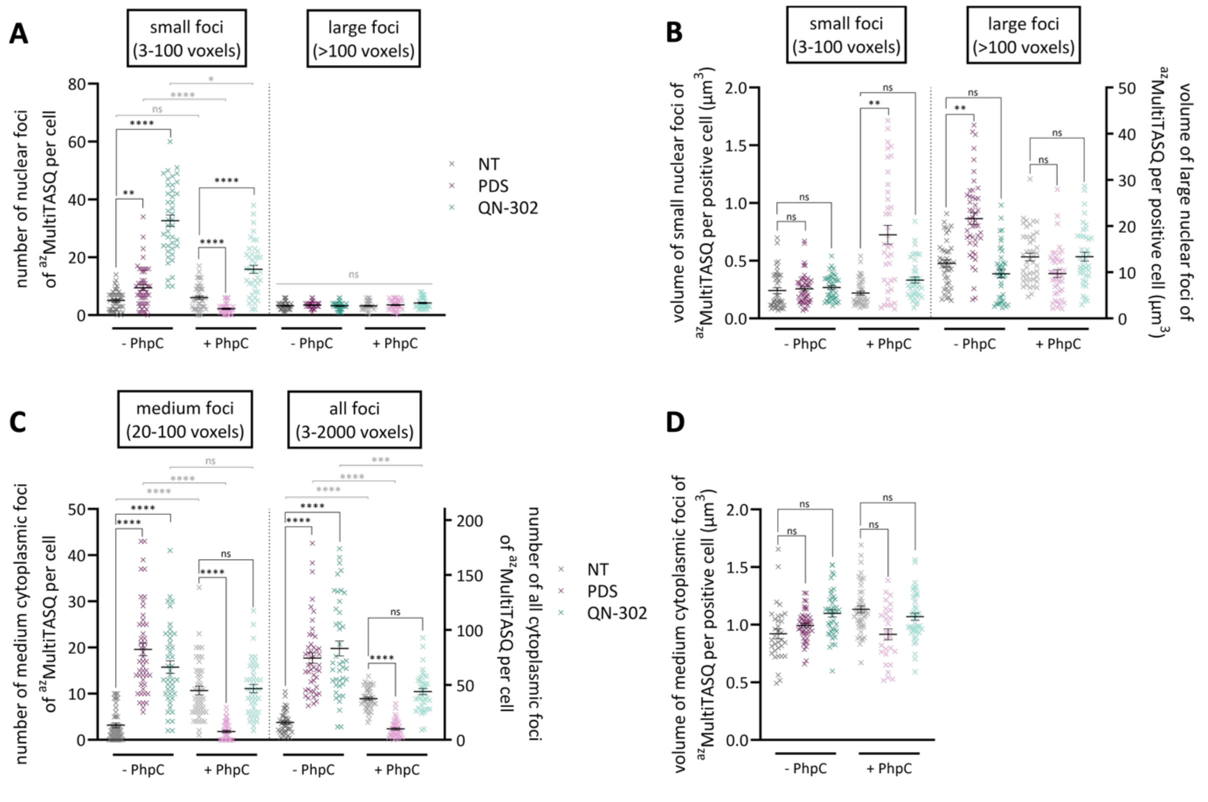


**Figure S3:** Quantification of $^{az}$MultiTASQ foci per cell in one independent experiment upon a 4-h G4-ligand treatment (PDS, 2 µM; or QN-302, 20 nM) with or without PhpC pre-incubation (20 µM). Quantification was performed from 50 ROI selected (as shown in **Fig. S2**) in cell image acquired with Olympus epi-fluorescence microscope equipped with 60x oil objective. **A.** Number of small (3-100 voxels) and large (>100 voxels; considered as nucleolar foci) nuclear TASQ foci quantified per cell. **B.** Mean volume (µm$^3$) of small (3-100 voxels) and large (>100 voxels; considered as nucleolar foci) nuclear TASQ foci quantified per cell, shown only in cells positive for nuclear TASQ foci (≥1 foci). Of note: we observed a significant increase of volume of small TASQ foci in +PhpC +PDS condition which could explain the significant decrease of number of small TASQ foci; however, the significant increase of volume of nucleoli upon -PhpC +PDS treatment is not correlated to a decrease of number of nucleoli and does not seem to impair small TASQ foci quantification still showing a correct segmentation of events of interest. **C.** Number of medium (20-100 voxels) and all-size (3-2000 voxels; obtained by adding the number of small, medium and large foci quantified for each ROI) cytoplasmic TASQ foci quantified per cell. Of note: we consider that medium foci quantified are representative of total number of foci without taking into account small foci that could be considered as background noise and large foci which could be poorly segmented and therefore give rise of incorrect number of foci. **D.** Mean volume (μm$^3$) of medium (20-100 voxels) cytoplasmic TASQ foci quantified per cell, shown only in cells positive for cytoplasmic TASQ foci (≥1 foci). Mean values and error bars (standard error of mean, sem) are represented as results collected from 50 ROI quantified per condition with outliers removal (n ≥ 42 per condition). Non-parametric one-way ANOVA with multiple comparisons (Kruskal-Wallis test) was employed for the statistical analyses with *: $P \leq 0.05$, **: $P \leq 0.01$, ***: $P \leq 0.001$; ****: $P \leq 0.0001$; “ns” stands for non-significant: $P > 0.05$.

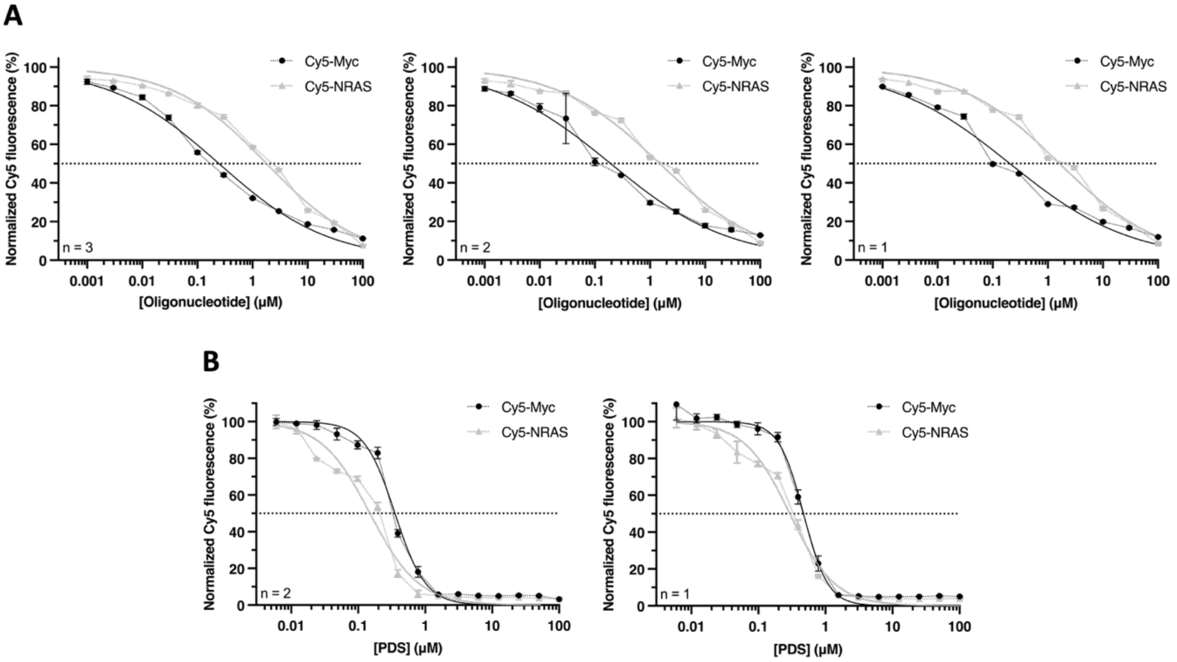


**Figure S4**: G4-affinity of QN-302 (200 nM) for either Cy5-Myc or Cy5-NRAS (from 0.001 μM to 1 mM) established by the reverse-FQA assay. G4-affinity of PDS (from 0.006 μM to 1 mM) for either Cy5-Myc or Cy5-NRAS (200 nM) established by the FQA assay. The experiments are performed at 25 °C in 50 mM Tris-HCl pH 7.2, 150 mM KCl, 0.05 % Triton X-100. The results are collected from triplicates (n = 3) across two (for PDS) or three (for QN-302) independent studies (n = 3).

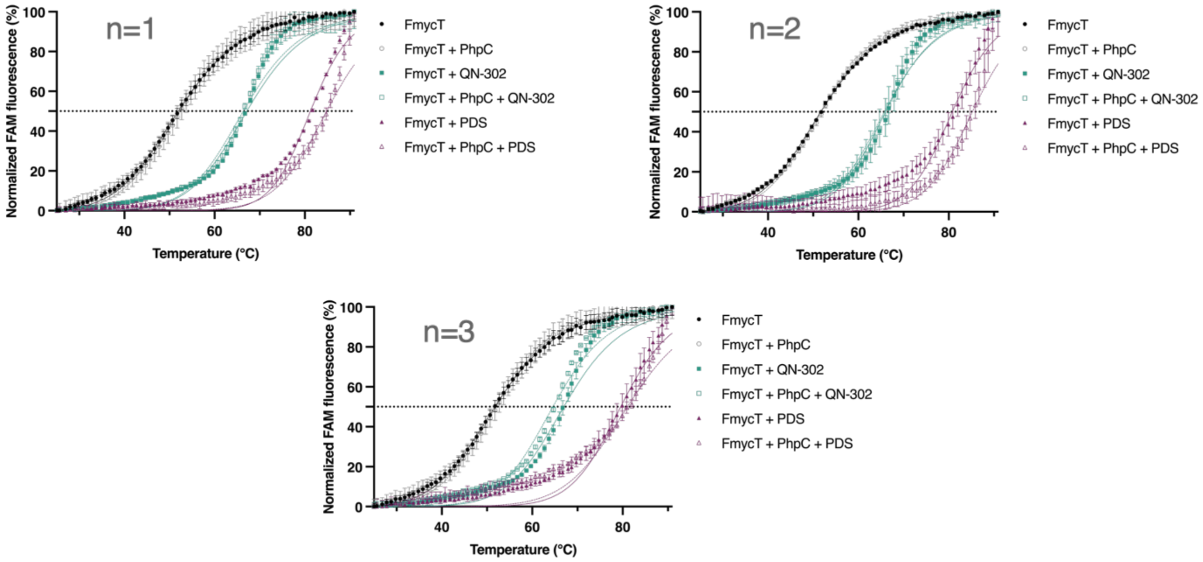


**Figure S5**: Competitive FRET-melting curves obtained with F-myc-T (0.2 μM) in presence of QN-302 (5 mol. equiv., 1 μM) or PDS (5. mol. equiv., 1 μM), without or with PhpC (50 mol. equiv., 10 μM). The experiments are performed in 10 mM lithium cacodylate pH 7.2, 1 mM KCl, 99 mM LiCl, from 25 to 90 °C

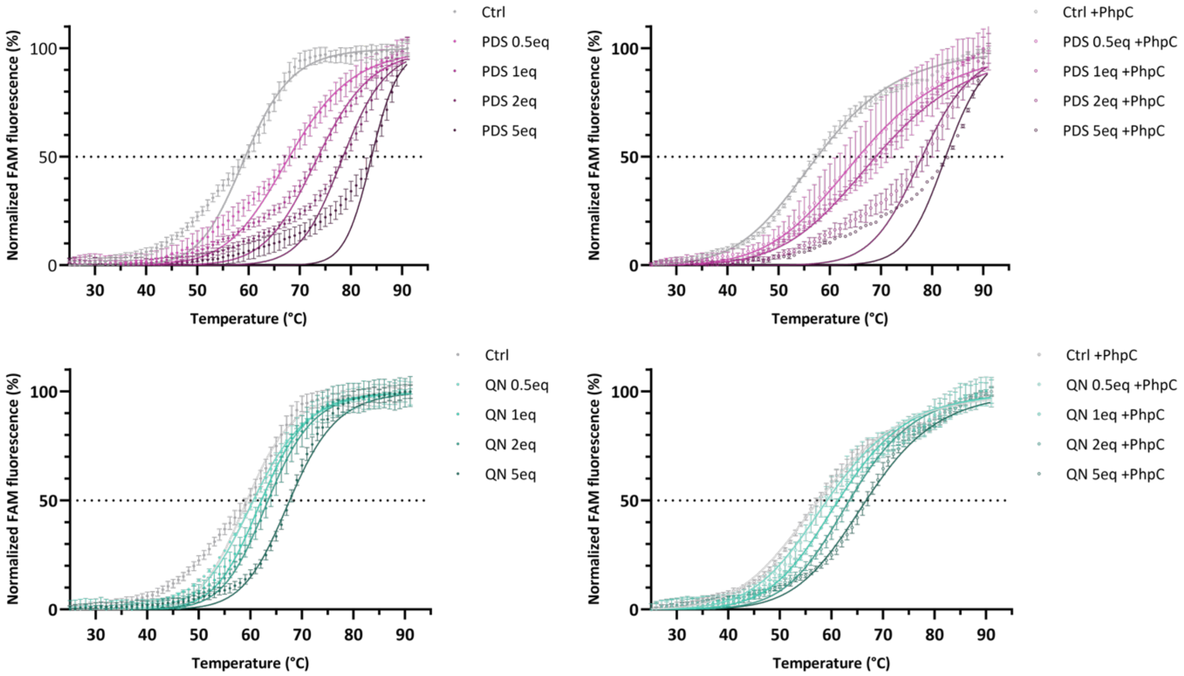


**Figure S6**: FRET-melting experiments performed with F-Myc-T (0.2 μM) and increasing concentrations of QN-302 and PDS (up to 1 μM) without (left) or with (right) pre-incubation (16 h at 4 °C) of PhpC (2 μM). The experiments are performed in 10 mM lithium cacodylate pH 7.2, 1 mM KCl, 99 mM LiCl, from 25 to 90 °C. The results are collected from a triplicate (n = 3), as a representative example.

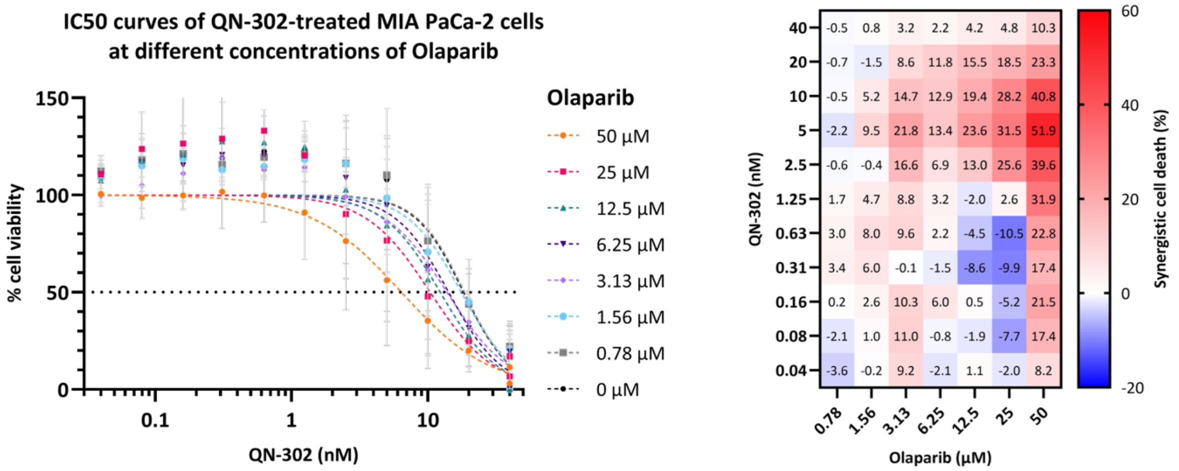


**Figure S7**: Synergistic interaction between QN-302 (up to 40 nM) and Olaparib (up to 50 μM) assessed by the SRB assay in MIA PaCa-2 cells after 72-h treatment. Cell viability curves were obtained after normalization to the untreated condition (without ligand) and then normalization of each QN-302 condition to the QN-302 untreated condition (left). Synergistic cell death (%) results in subtraction of each cell viability value to drug untreated condition (right). The results are collected from three independent studies (n = 3).

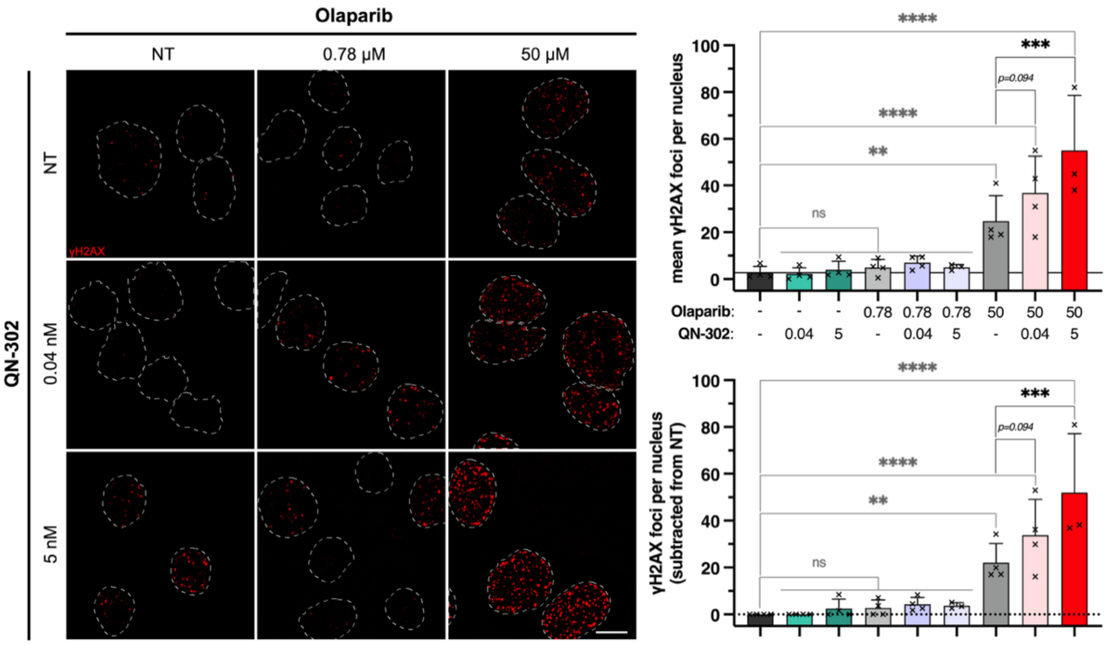


**Figure S8**: Synergistic interaction between QN-302 (0.04 or 5 nM) and Olaparib (0.78 or 50 μM) assessed by the immunodetection of DSBs with a combination of four high/low concentrations after a 24-h treatment in MIA PaCa-2 cells. Representative image (left) of observed synergistic effect quantified by γH2AX *foci*/cell (right). These results are collected as triplicates. Fisher's LSD tests were employed for the statistical analyses with *: P ≤ 0.05, **: P ≤ 0.01, ***: P ≤ 0.001; ****: P ≤ 0.0001; "ns" for non-significant: P > 0.05.

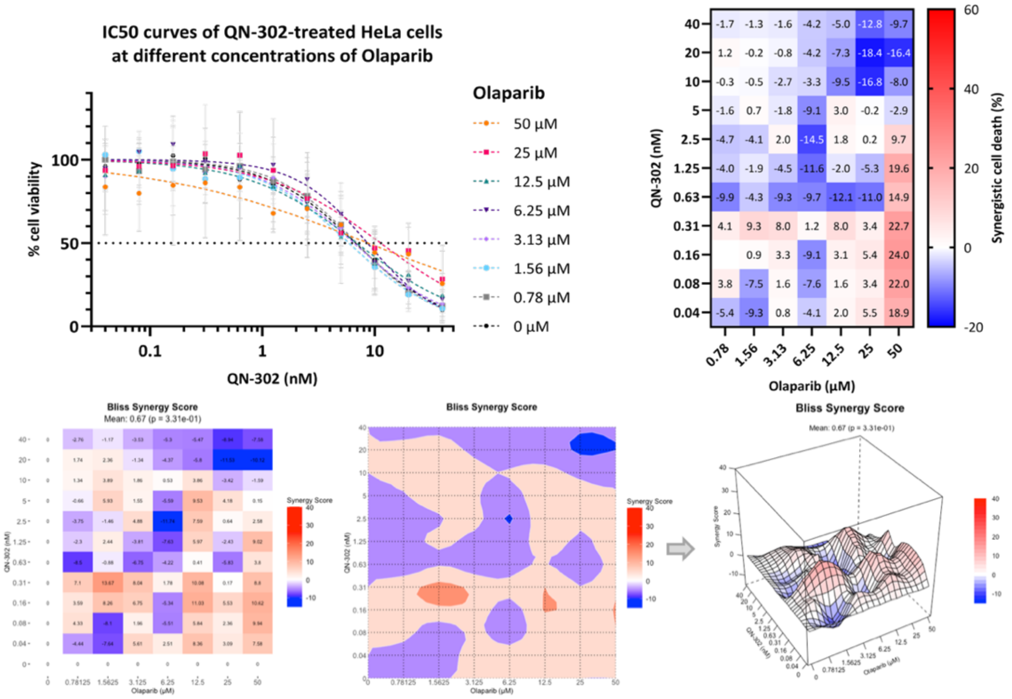


**Figure S9**: Synergistic interaction between QN-302 (up to 40 nM) and Olaparib (up to 50 µM) assessed by the SRB assay in HeLa cells after 72-h treatment. Cell viability curves were obtained after normalization to the untreated condition (without ligand) and then normalization of each QN-302 condition to the QN-302 untreated condition (left). Synergistic cell death (%) results in subtraction of each cell viability value to drug untreated condition (right). The results are collected from three independent studies.

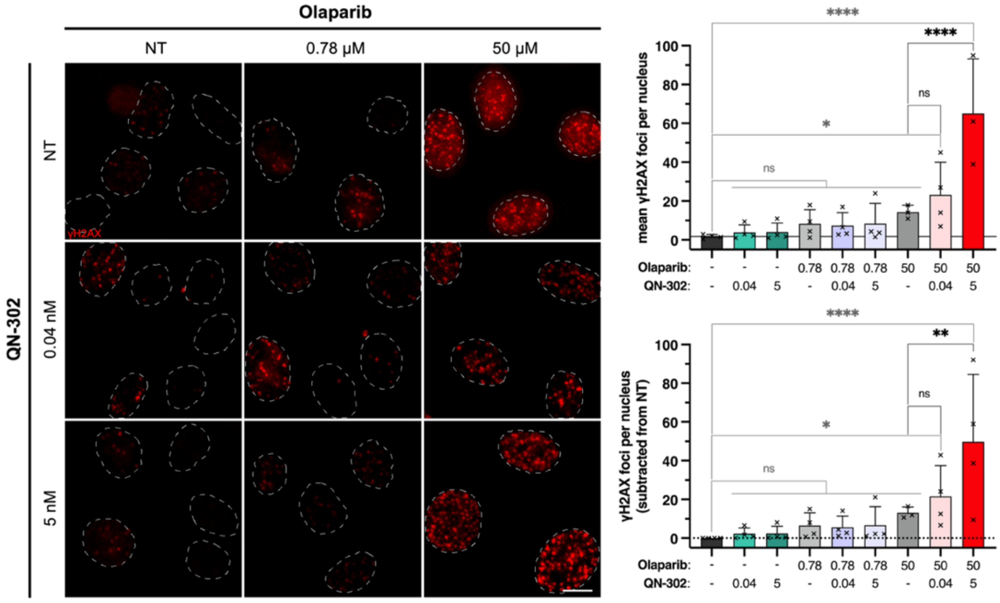


**Figure S10**: Synergistic interaction between QN-302 (0.04 or 5 nM) and Olaparib (0.78 or 50 µM) assessed by the immunodetection of DSBs with a combination of four high/low concentrations after a 24-h treatment in HeLa cells. Representative image (left) of observed synergistic effect quantified by γH2AX *foci*/cell (right). These results are collected as triplicates.. Fisher's LSD tests were employed for the statistical analyses with *: P ≤ 0.05, **: P ≤ 0.01, ***: P ≤ 0.001; ****: P ≤ 0.0001; "ns" for non-significant: P > 0.05.
